# Dimensionality and the statistical power of multivariate genome-wide association studies

**DOI:** 10.1101/016592

**Authors:** Eladio J. Márquez, David Houle

## Abstract

Mutations virtually always have pleiotropic effects, yet most genome-wide association studies (GWAS) analyze effects one trait at a time. In order to investigate the performance of a multivariate approach to GWAS, we simulated scenarios where variation in a *d-*dimensional phenotype space was caused by a known subset of SNPs. Multivariate analyses of variance were then carried out on *k* traits, where *k* could be less than, greater than or equal to *d*. Our results show that power is maximized and false discovery rate (FDR) minimized when the number of traits analyzed, *k*, matches the true dimensionality of the phenotype being analyzed, *d*. When true dimensionality is high, the power of a single univariate analysis can be an order of magnitude less than the *k=d* case, even when the single trait with the largest genetic variance is chosen for analysis. When traits are added to a study in order of their independent genetic variation, the gains in power from increasing *k* up to *d* are much larger than the loss in power when *k* exceeds *d*. Simulations that explicitly model linkage disequilibrium (LD) indicate that when SNPs in disequilibrium are subjected to multivariate analysis, the magnitude of the apparent effect induced onto null SNPs by SNPs carrying a true effect weakens as *k* approaches *d*, such that the rank of P-values among a set of correlated SNPs becomes an increasingly reliable predictor of true positives. Multivariate GWAS outperform univariate ones under a wide range of conditions, and should become the standard in studies of the inheritance of complex phenotypes.

## Introduction

The vast majority of genetic association studies focus on one trait at a time. Even when more than one trait has been measured, separate analyses are usually conducted for each trait. For example, van der Harst [1] characterized a human population for six related red blood cell phenotypes by testing for association in each phenotype independently. Even studies of transcriptome abundances that sample thousands of RNAs [2] usually analyze them individually. In other cases, the dimension of multivariate data is first reduced by principal components analysis, and scores on the trait combinations that define a PC are then analyzed independently [e.g., 3]. There is increasing interest in inferring joint effects on more than one trait to characterize pleiotropy [4–6], although in many cases this has been accomplished by reanalysis of the results of univariate analyses [e.g., 7,8,9]. Such analyses are actually biased against discovering pleiotropy, as the associations must meet strict univariate criteria for significance before being considered for testing [6].

A truly multivariate analysis, where a single data set is analyzed simultaneously for the relationship between *m* genetic markers (e.g., single nucleotide polymorphisms, SNPs) and *k* traits, offers several advantages over such reanalyses of univariate associations. First, it is efficient in the number of tests, as each SNP is subjected to a single analysis, rather than *k* analyses. Second, multivariate tests are more powerful than sets of univariate tests as they locate those directions of phenotype space that maximize phenotypic differences among predictors, regardless of their orientation relative to the phenotypic measurements. Third, multivariate analyses estimate the direction of effects in phenotype space. This enables quantitative interpretation of the sources of covariation among the measured variables. It also provides a quantitative basis for validation of results in later studies.

While a small number of such truly multivariate genome-wide analyses have been published [e.g., 10,11], the potential advantages of such analyses have received relatively little attention. Statisticians have adapted a variety of multivariate statistical approaches to multivariate association studies [6,12,13], but with few exceptions [e.g., 14,15], most have tended to focus on problematic aspects of large data sets, such as dealing with missing values, generalizing to non-normally distributed data, or including mixtures of continuous and categorical phenotypes, than on the performance of simple multivariate analyses.

The opportunity for such multivariate analyses is certain to grow for several reasons. First of all, there is recognition that understanding the genotype-phenotype map is an inherently multivariate problem. The totality of genetic effects on the phenotype as a whole is what shapes its evolutionary, medical and economic significance. This has led to increasing emphasis on phenomics, high-throughput acquisition of multivariate phenotypic data to complement our ability to measure genomes [16]. Second, there are now many efforts underway to generate and phenotypically characterize replicable sets of genotypes in a variety of model organisms [e.g., 17,18,19], greatly increasing the number of datasets where a truly multivariate analysis can be undertaken. Complementary efforts in humans aim to increase the comprehensiveness of phenotyping efforts [e.g., 20,21] and to increase the sizes of cohorts studied [e.g., 22].

Our purpose in this contribution is to explore the statistical properties of multivariate genome-wide association mapping through simulation when the true dimensionality of genetic effects is not known. To understand our approach, we need to define our use of the term dimensionality. Dimensionality is the number of independent *aspects of the phenotype* that are affected by genetic variation in the suite of traits measured. Independence is defined geometrically as orthogonal directions in the space of all measured traits. For example, imagine that the measured traits are the length and width of some morphological structure. Clearly if genetic variation in the population affects only length or only width, then the true dimension of variation (*d*) is one. Critically, if all genetic changes affected length and width in the same way – say changing length twice the amount that width is changed – *d* remains one, even though two traits are affected by each mutation. When different genetic variants affect length and width in different ways, then the true dimension of variation is at least *d*=2. While the true dimensionality can be less than the number of traits measured, we have to measure at least *d* traits to test the hypothesis that the dimensionality of variation is at least *d*. Critically, length and width can be correlated, yet measuring both adds information that allows that second orthogonal dimension of variation to be assessed. In general, each of the *d* dimensions will be defined by some combination of the characteristics measured. Our definition of dimensionality generalizes more conventional usages such as “correlated traits,” as it emphasizes conditionally independent directions of genetic variation, over specific measurements (such as length and width), which an investigator chooses to measure.

It is clear that multivariate analysis will be greatly superior when genetic variants can have any possible combination of effects on the phenotype [14,15], that is, when the dimensionality is as large as the number of measured phenotypes, and includes all possible direction in the space of measured phenotypes. However, it is possible to measure more phenotypes than the true genetic dimensionality, and thus include phenotypic measurements that do not contribute independent information about genetic causation. Consequently, we need to consider not just the benefits of measuring more traits when the true dimensionality is high, but also the costs of analyzing too many phenotypes when the true dimensionality is low. We restrict our attention to the case of multivariate normal phenotype distributions that can be analyzed by standard multivariate analyses of variance (MANOVA).

We present results of two sets of simulations to address the properties of MANOVA of high dimensional data for association mapping. Our first set of simulations examines the effect of trait dimensionality on statistical performance of single-locus tests for the ideal case where linkage (gametic) disequilibrium (LD) is absent. This focuses attention on how multivariate analyses might be different from univariate ones. Second, we explore the statistical consequences of LD by simulating phenotypic variation caused by one of a set of correlated SNPs. Our findings display the widely reported statistical challenges posed by genome-wide analyses in general, but clearly demonstrate that increasing the number of phenotypes usually leads to an increase in statistical power.

## Simulation Methods

The primary aim of these simulations is to create *k*-dimensional phenotypic data with known causal associations to a SNP data set. We consider the simple case of a set of traits that follow a multivariate normal distribution, so that a standard multivariate analysis of variance is appropriate. The parameters of our simulations were based on the properties of the Drosophila Genome Reference Panel [DGRP; 19], a set of inbred lines derived from a single outbred population of *Drosophila melanogaster*. This population has been used for both published and ongoing association mapping efforts.

### Genotype information

We used Freeze 1 genotype calls (ftp://ftp.hgsc.bcm.edu/DGRP/freeze1_July_2010/snp_calls/) for a subset of 164 DGRP lines as the basis for our simulations. Details of the SNP calling can be found in Mackay et al. (2012). We used only calls of homozygous genotypes at sites with exactly two alternative types. All other calls were treated as missing data. There were 2,489,796 sequenced sites where the count of the minor allele was at least 3, and the number of non-missing calls was at least 61. We further filtered these sites for local gametic (linkage) disequilibrium by calculating the correlation of genotypic identities between all sites within 20 polymorphic sites along each chromosome. For pairs of sites with *r*^2^ > 0.8, we dropped the site with the greater number of missing calls, and when these were the same, dropped the site with the lowest minor allele count. In practice, every dropped site proved to be perfectly correlated (*r*^2^ = 1) with at least one included site. After this site selection step, 1,541,611 sites remained. The median allele frequency was p=0.121, and the median number of lines scored at each SNP was 160.

### Simulating phenotypic effects in the absence of LD

For the simulations constructed without LD, we generated *m* = 10,000 random arrays of allelic values whose minor allele counts (MAC) and number of lines with missing genotype data were sampled randomly from the Freeze 1 DGRP sites. To ensure that LD was random, we randomly assigned SNP genotypes and missing values to lines. A subset of SNPs were randomly chosen to cause simulated effects, and thus contribute to the genetic variance among simulated genotypes. Following Storey and Tibishirani [23], we denote the fraction of loci that do *not* have an effect as π_0_, so that the total number of sites with a real effect is *m*_1_ = *m* (1- π_0_).

The directions of SNP effects were based on the among-line variance-covariance matrix **G**, derived from the positions of 48 landmark and semi-landmark points on the wings of female specimens from 164 DGRP lines (N=3,635). We defined our traits to be the first 59 non-null eigenvectors (the theoretical maximum number of independent traits in our data), resulting in a diagonal covariance matrix **Ĝ**. Thus, our simulated multivariate traits have simple factor structure, with each ordered dimension capturing a progressively smaller proportion of the total genetic variance. We assumed that a proportion *u* of the genetic effects captured in **Ĝ** were not caused by any of the SNPs in our sample, while the remaining 1-*u* was. For these simulations, we held *u* constant at 0.1. We then partitioned **Ĝ** into two orthogonal complementary subspaces, namely **L** = (1 − *u*)**Ĝ**, which parameterizes the contribution due to known SNPs, and **L**_**res**_ = *u***Ĝ**, which models other genotypic effects.

To generate SNP effects, we randomly sampled *m*_1_ multivariate normal effect vectors, *S*_*j*_ with mean vector **0** and covariance parameter matrix **L**. We then normalized the *S*_*j*_ vectors and assigned each a magnitude *g*_*j*_, sampled from an exponential distribution with mean 1. The total effects of all SNPs in line *i* is then Σ_*j*_ *a*_*ij*_*g*_*j*_*S*_*j*_, where *a*_*ij*_ denotes the allele for genotype *i* at locus *j*, valued 0 if it corresponds to the major allele and 1 if it equals the minor allele. For the purposes of computing genotype means, we imputed missing alleles a random value (0 or 1). To maintain comparability across simulations, we scaled the effect size parameter *g* so that the trace of the simulated covariance matrix matched the trace of **L**, using,

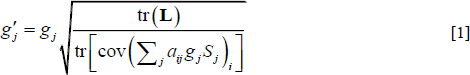

where tr and cov denote the trace and covariance matrix operators. Constraining simulated covariance matrices to account for a constant variance holds the proportion of variation accounted for by all SNPs constant, regardless of the number of sites causing that variation. This means that the proportion of variance explained by each SNP is larger in simulations with fewer causal SNPs. We ran simulations based on high- and low-power scenarios, assuming, respectively, *m*_1_ =100 (π_0_ = 0.99) and *m*_1_ = 3000 (π_0_ = 0.70) causal SNPs. Figures S1 and S2 show, respectively, typical distributions of effect sizes and per SNP explained variance simulated under these power scenarios.

To generate unexplained genetic differences among lines, we randomly sampled a vector, *U*_*i*_, from a multivariate normal distribution with mean vector **0** and covariance matrix **L_res_** for each of the *N* inbred genotypes. We drew samples of *U*_*i*_ repeatedly until we obtained a set where the covariance matrix of these effects, **U**, satisfied tr(**L_res_**)-tr(**U**) < ε. We set ε=10^−6^.

The mean phenotype of the *i*th line, *Y*_*i*_, is the sum of all these genetic effects

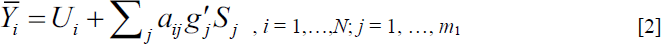

Finally, we simulated samples of *n* = 40 individuals within genotypes by drawing random vectors from a multivariate normal distribution with mean 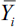 and covariance matrix **E** computed by subtracting **G** from the pooled within-line covariance matrix (**P**). The structure of the **L** and **E** matrices parameterized from DGRP data have the relatively even eigenvalue distributions expected for morphometric data [16]. The maximum eigenvalues of **L** and **E** are 1.18 and 0.33, respectively, and the slopes from regressing logarithm of eigenvalues on *d* are *β*_L_ = -0.07 and *β*_E_ = -0.05. Each successive eigenvalue is on average 85% of the previous eigenvalue for the among-line variation, and 89% of the previous eigenvalue in the unexplained matrix. Consequently, genetic variation drops more sharply at higher dimensions than unexplained variation does, resulting in a steady decline of heritability (*h*^2^) with dimensionality (see Fig. S4).

### Simulating phenotypic effects with disequilibrium

To include the effect of LD in simulations, we designed a sampling scheme that would allow us to isolate the biasing impact of correlations among SNPs on the probability that a null SNP is declared significant (i.e., false positives) by virtue of its correlation to a true SNP, whether the latter is declared significant (i.e., true positive) or not (i.e., false negative). To generate correlated SNPs, we drew samples of SNP genotypes directly from the filtered Freeze 1 DGRP sequences. To ensure an even representation of allelic frequencies, we ranked all sites by their minor allele frequency (MAF), and then sampled focal SNPs from within each MAF decile at random. We identified all SNPs genome-wide whose gametic correlation with each focal SNP satisfies *r* > 0.5, and denote the resulting set as a “SNP family.” We retained for analysis 100 families with at least two members (the focal SNP and at least one other) from each MAF decile. The mean and median number of SNPs/family was 169 and 14 respectively. In cases where the focal SNP was correlated at *r* > 0.5 with more than 100 SNPs (31% of all focal SNPs), we retained a random sample of 100 correlated SNPs.

To isolate the impact of LD in the presence of an effect, we carried out paired simulations where either the focal SNP in a family was assigned a phenotypic effect, or none of the SNPs in the family had phenotypic effects. In these simulations each effect explains 1% of the among-line variance, with the remainder left unexplained (*u* = 0.99). Simulations were otherwise performed as described in the absence of LD. This allowed us to precisely quantify the statistical impact of a genetic effect in a sample of correlated SNPs.

### Dimensionality

True dimensionality, *d*, is the number of dimensions of the space in which genetic effects on the phenotype arise. We wanted to explore the implications of the number of measurements analyzed, *k*, in relation to *d*. Our diagonal covariance parameter matrix **Ĝ** spans the entire 59-dimensional space of possible genetic variation. To generate a *d*-dimensional parameter covariance matrix **Ĝ**_*d*_, we simply set the bottom 59-*d* diagonal elements of **Ĝ** to 0. Thus, the *d*-dimensional subspace retained is always the subspace that contains the most phenotypic variance. This procedure is conservative with respect to the usefulness of a multivariate approach, because the first dimension always captures more of the genetic variance than any other dimension, and our univariate analyses are of this one most informative trait. Finally, to avoid confounding differences in dimensionality and total variance across simulations, the total variance captured in the first *d* dimensions is rescaled to match the trace of **Ĝ**, by multiplying each element of **Ĝ**_*d*_ by (**Ĝ**)/tr(**Ĝ**_*d*_). Note that phenotypic data are in all cases measured in the original 59-dimensional space, where the bottom 59-*d* dimensions contain non-genetic residual variation, as described by **E**.

### Analysis

We used MANOVA (for *k* > 1 simulations) and ANOVA (for *k* = 1 simulations) to test for genomic associations, with SNP genotype as the sole predictor variable and *k* (= 1, 2, 5, 10, 20, 30, 40, 50, and 59) shape traits as response variables. MANOVA P-values were calculated using a chi-square approximation of Wilk’s Lambda, whereas *F*-ratios were used for ANOVAs. We also ran an additional set of simulations for the case where *d*=59 based on *k* univariate tests. In these tests, a SNP was declared significant if any of the *k* P-values were less than the critical value. Critical values for these simulations were Bonferroni-adjusted as α* = α/*k*.

For each combination of the parameters *k, d*, and π0, we generated 100 replicate samples of *m*_1_ effect SNPs. To generate Type I error and FDR estimates, we drew a random sample of *m*- *m*_1_ P-values for each replicate set of effect SNPs from a pool of ~10^6^ P-values computed from null SNPs in preliminary simulations. This simplifying procedure is based on the well-known expectation that P-values from null SNPs follow a uniform distribution (e.g., Hung et al., 1997), and it allowed us to increase replication of the simulation study while minimizing computer run times. We validated our results by running two fully sampled replicates where *m* = 10,000 SNPs were simulated at both power scenarios using all *k* and *d* combinations shown here.

Simulations of SNP and phenotypic data were carried out in Matlab r2010b [24]. Simulated data were analyzed using SAS/STAT PROC GLM software, Version 9.2 of the SAS System for Windows and Unix. A software package to carry out these simulations has been made publicly available at http://bio.fsu.edu [Note to editor: URL is temporary; software will be made available upon publication]. Code is available upon request from the corresponding author.

## Results

### Independent SNPs

The simulations are constructed so that we know which predictor SNPs are assigned effects, which we term “effect SNPs,” and the “null SNPs” that do not have effects. Once our simulated data have been subjected to statistical testing, the *m* SNPs are categorized into four groups as shown in Table 1. This allows us to understand the relationship between the parameters of the simulations and tests and the following measures: Power, or true positive rate (TPR=TP/[TP+FN]), is the proportion of effect SNPs that are declared significant. False positive rate (FPR=FP/[FP+TN]) is the proportion of null SNPs that are declared to be significant. False discovery rate (FDR=FP/[TP+FP]) is the proportion of SNPs judged significant that are null. If the assumptions of the statistical test are met, Type I error is equal to the critical P-value, α. Type II error, *β*, is determined by properties of the data and the choice of α. Type I and Type II error rates determine power, FPR and FDR.

**Table 1.** Number of test outcomes categorized according to the presence of a true effect and the outcome of the test.

|  | Test result |  |  |
| --- | --- | --- | --- |
| SNP | Significant | Non-significant | Total |
| Effect | $TP = m(1-\pi_0)(1-\beta)$ | $FN = m(1-\pi_0)\beta$ | $TP+FN = m(1-\pi_0)$ |
| Null | $FP = m\pi_0\alpha$ | $TN = m\pi_0(1-\alpha)$ | $FP+TN = m\pi_0$ |

TN: true negative count, FP: false positives, FN: false negatives, $\pi_0$ :
proportion of tests for which the null hypothesis of absence of effect is true, $\alpha$ and $\beta$ : Type I and
II error, respectively, $m$ : total count of tested hypotheses.

The independent SNP simulations are constructed to match the assumptions of our statistical tests. We tested for departures from the expected uniform distribution of P-values computed from null SNPs, and verified that the FPR matched expected Type I error rates in the absence of true effects. In all cases, results conformed to the null expectation.

### Power and FDR

The first set of simulations is directed at understanding the relationship between power, the number of dimensions that have genetic effects (*d*), and the number of orthogonal measurements analyzed (*k*). In each replicate, we sampled *m*=10,000 independent SNPs, and assigned phenotypic effects to *m*_*1*_ = *m*(1-π_0_) of these as described in the Methods. The *m*_*1*_ SNPs were assigned effects sufficient to explain 90% of the genetic variation observed in our experiments (i.e., *u* = 0.1). We simulated data for nine different values of *d*; each of these sets was then analyzed at the same nine values of *k*. Each (*d, k*) combination was replicated 100 times. We show results for a high-power case where 100 of the SNPs cause the genetic variance in phenotype (π_0_=0.99), and a low-power case where 3,000 of the SNPs cause the genetic variance in phenotype (π_0_=0.70).

Figures 1 and 2 show power and FDR for a variety of α values for the high and low power simulations. Power is, as expected, strongly influenced by 1-π_0_, the proportion of SNPs with phenotypic effects. Power is much higher overall when the genetic variance can be explained by a smaller number of SNPs of large effect. Similarly, power can be increased by relaxing α, although this has the expected [25] and detrimental effect of making the probability that a SNP is judged significant essentially random, so that FDR approaches π_0_. The underlying exponential distribution of effect sizes simulated is a challenging one for GWAS, as the mode of effect size will be close to 0. When *k* and *d* are both high, power can be impressively high, given this distribution of effect sizes.

**Figure 1.**
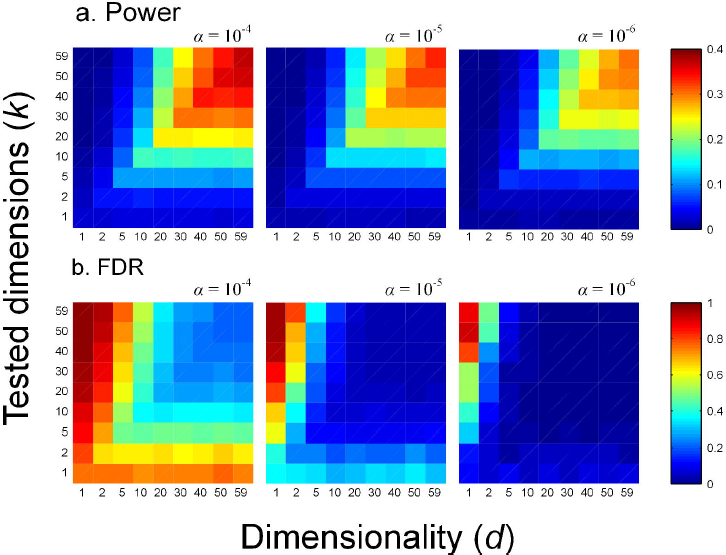
Power (a) and FDR (b) simulated as a function of dimensionality (*d*) and number of tested dimensions (*k*) for the high-power scenario.

**Figure 2.**
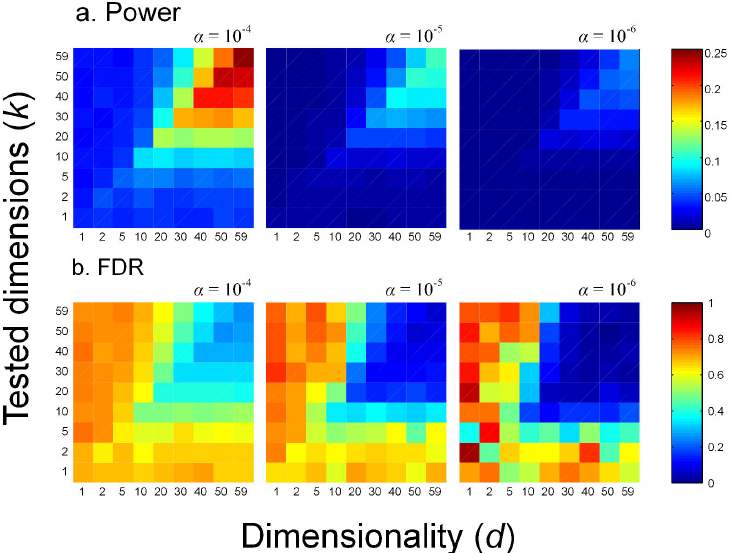
Power (a) and FDR (b) simulated as a function of dimensionality (*d*) and number of tested dimensions (*k*) for the low-power scenario.

The key results relevant to whether a multivariate approach to GWAS is beneficial are in the pattern of power and FDR at different dimensionalities (*d*) and numbers of traits (*k*). Despite the obvious wide differences that varying power and α have, the pattern of effects of *d* and *k* are similar across the range of parameter values. These patterns are shown most clearly in Figure 3, which gives a more detailed look at power and FDR for the high-power case at α = 10^−4^. First, note that when *k* is low, power is always low and FDR is always high. This is most evident in the univariate case, utilized in almost all previous association studies. When *d* is low, performance is also poor, regardless of *k*. Second, power increases and FDR decreases with *k* when *k*<*d*. Power is maximized and FDR minimized when the true and tested dimensionalities match, that is when *k*=*d*. Third, power decreases and FDR increases modestly when *k*>*d*, showing a diminishing rate of change as *k* increases. At *k*<*d*, increasing *k* places the true effects against a smaller background of error variance in the direction of the effect. When *k*>*d*, increasing *k* increases the error variance, while leaving the signal of the true effects unchanged. Clearly, when the underlying genetics is multivariate, a multivariate analysis is superior to a univariate analysis.

**Figure 3.**
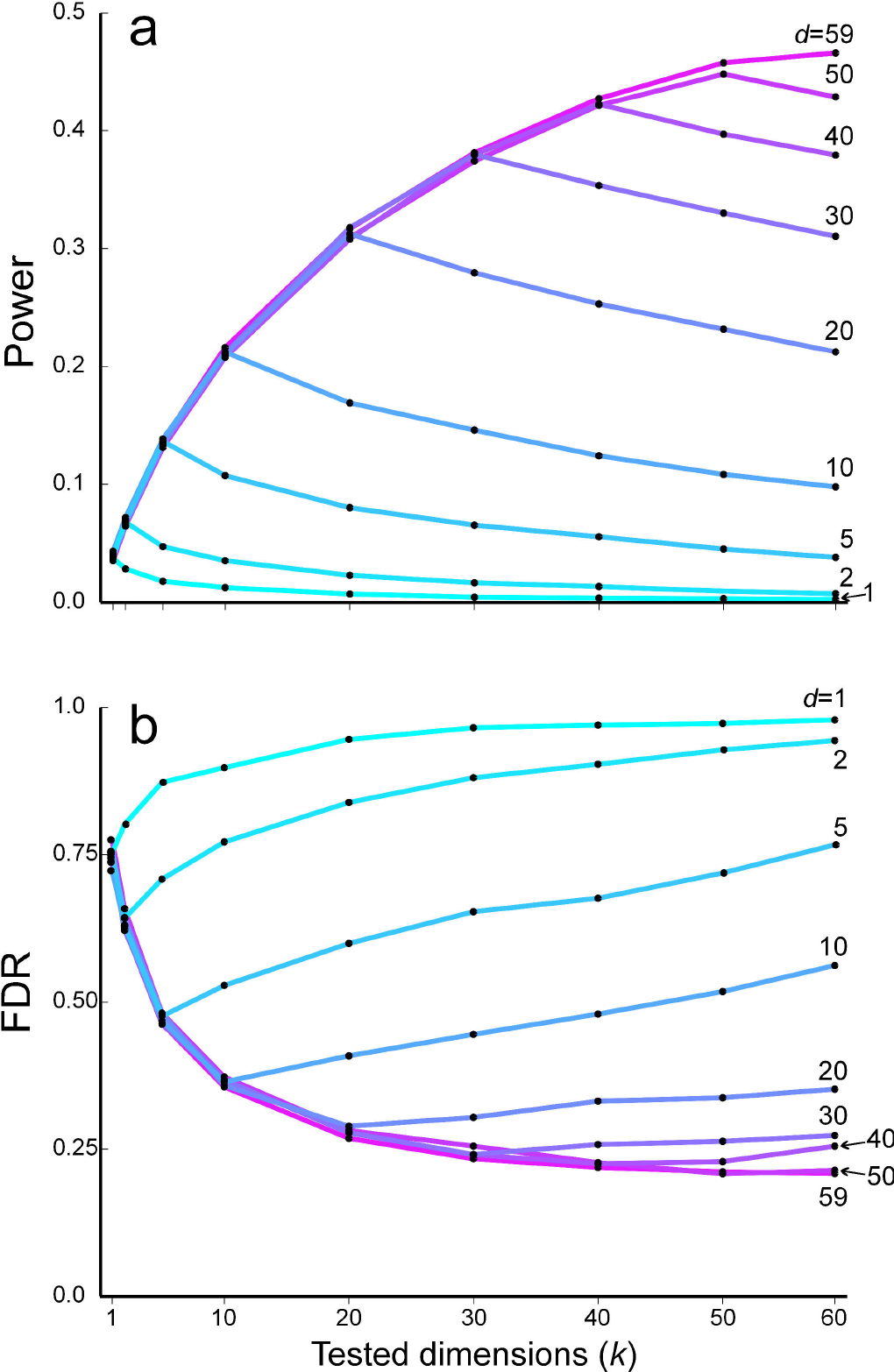
Effect of number of tested traits (*k*) on (a) power and (b) FDR for various dimensionalities (α=10^−4^). Data correspond to the high-power simulations (π0=0.99) shown in Figure 1.

The asymmetry around *k*=*d* is likely due to our choice of variance-ordered principal components scores as traits, so that trait *i* is always more variable than trait *i*+1. In other words, we always choose the most informative *k* traits to measure. Thus patterns shown in Figs. 1–3 should hold when trait sampling is based on preliminary data that identify those aspects of the phenotype that are most informative, rather than a random subset of possible traits. If we had not chosen to treat principal component eigenvectors as traits, the variance explained by adding one more trait would have a far greater random component, although the expected gain (or loss) will be the same. In some cases the *k*+1*th* random trait would explain more conditionally independent variation than the average of the first *k* random traits, but in other cases, it will explain less. Our results reflect the average gain over adding all possible *k*+1*th* traits that define a particular phenotype space.

When confronted with a multivariate data set with *k* variables, a common response is to perform *k* univariate analyses. In Figure 4 we compare the SNP-wise power and FDR from such multiple univariate analyses to the corresponding *k*-variate multivariate analyses using the *d*=59, high-power simulations whose results are shown in Figures 1 and 3. To perform the multiple univariate analysis, we calculated the probability that a SNP was declared significant in any of the *k* univariate tests after applying a Bonferroni correction to α. The power of the multiple univariate analysis relative to the corresponding multivariate analysis decreases as *k* increases. Similarly the relative FDR increases. On the other hand, the *k* univariate analyses are far more effective than a single univariate analysis at detecting SNP effects. For example, examination of Fig. 3A shows that at α = 10^−4^ a 59-dimensional multivariate analysis is about 10 times more powerful than a single univariate analysis, whereas Fig. 4A (circles) shows that the same analysis is only 2.2 times as powerful as 59 univariate analyses. Similarly, the lower P-value used in the multiple-univariate analyses is effective at controlling the FDR rate relative to single univariate tests.

**Figure 4.**
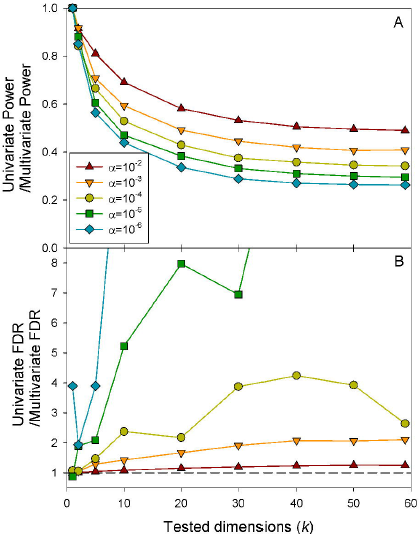
Power (a) and FDR (b) of *k* univariate tests relative to a single *k*-dimensional multivariate test.

Additional simulations show that the uniformly low power for univariate analyses in our simulations can be partially rescued by increasing the number of lines included in the study. For example, for the high-power scenario described here (π_0_ = 0.99), using α = 10^−4^, a five-fold increase in sample size from *N* = 164 to 820 results in a five-fold increase in average power from 2.3% to 10.5% for the univariate case (*k* = 1) when *d* = 59, whereas for the corresponding *k* = *d* = 59 case, average power increases from 41.8% to 69.8%. Power also increases modestly when SNP effect sizes are more homogeneously distributed than they are under the exponential distribution, or when the number of causal SNPs is even smaller (results not shown).

### Simulations of correlated SNPs

The above simulations were constructed to ensure that the SNP genotypes are independent, that is that there is no LD. This is clearly unrealistic, as any genome-wide association study will include closely linked SNPs that may be in LD, as well as distant sites that happen to be correlated in the finite sample of genotypes scored. Consequently, we conducted an additional set of simulations focused on the effects of LD on power and FDR. As described in Methods, we used a real data set, the Drosophila Genome Reference Panel [19], to provide an example of LD in a real population to structure our simulations. From these data, we identified all SNPs correlated with a focal SNP, which we call a SNP family. We simulated effects for focal SNPs in 1,000 SNP families, 100 families drawn from each MAF decile that had at least one correlated SNP. The mean and median number of SNPs/family was 169 and 14, respectively.

Interpretation of results in the presence of LD is more complex than the ideal case, and depends on the purpose of the analysis. If the goal is to confirm that some genetic variation affects the phenotype, then any significant SNPs in a family with a true positive SNP is helpful. We refer to such correlated SNPs as family-positives. We would ideally like to identify individual SNPs that cause phenotypic variation (QTNs), so that, for example, the nature of regions with effects can be studied. In this case, we seek the true-positive causal SNP.

Figure 5 shows the relevant results for the case of *d*=59, *k*=59. Regardless of the P-value cutoff, the number of family-positive SNPs per true positive SNP (upward pointing triangles) remains on average above 1. Note that even when the focal SNP with the simulated effect is not significant (downward pointing triangles in Fig. 4), there are frequently (but not always) correlated family-positive SNPs. This suggests that in the presence of even the modest disequilibrium present in the DGRP, the SNP-wise false discovery rate will be high. Note that the SNP-wise false positive rate in the absence of a simulated effect was always higher than the nominal value due to the large number of SNPs in the average family. For the effect size simulated, power remains good over the entire range of P-values.

**Figure 5.**
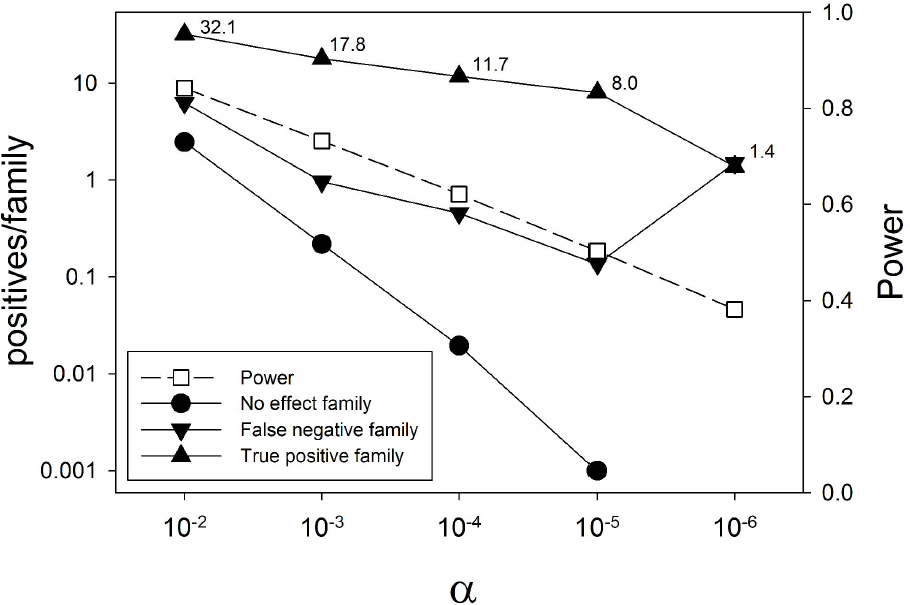
Base-10 logs of mean counts of family-positives per family of correlated (*r* > 0.5) SNPs as a function of the significance cutoff α. This shows the bias of P-value distributions for SNPs in LD with a SNP that carries an effect. We contrast cases where the effect SNP reaches significance (true positive; upward-pointing triangles) with those where the effect SNP is non-significant (false negative, downward pointing triangles). Also shown is the case where no SNP in the family has an effect (circles: all significant SNPs are false positives). Open squares denote power to detect true effects (i.e., true positive rate). No family positives in true-effect families were observed at P<10^−6^.

We investigated how the family-positive rate changes with the magnitude of LD. Figure 6a shows the probability that a null SNP is significant when the focal SNP is correctly judged significant, as a function of gametic correlation (*r*) with the focal SNP. The family-positive rate per SNP declines rapidly as *r* decreases. On the other hand, Fig. 6b shows that this decrease in probability of a family-positive per SNP is countered by the much larger number of SNPs with weaker correlations (Fig. 6c). When less conservative values for α are used, the number of family-positives correlated with each focal SNP continues to rise as lower *r* bins are considered. Even when conservative P-values are adopted, the advantage suggested in Fig. 6a is much reduced. Note that sites perfectly correlated with the focal SNP can still yield different test statistics than the focal SNP due to missing genotypic assignments.

**Figure 6.**
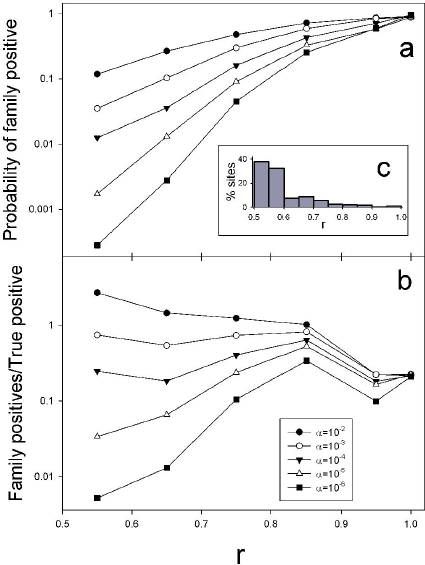
Family positives as a function of gametic correlation. (a) Probability that a null SNP correlated with a true positive SNP tests positive for sites binned by gametic correlation (*r*), for five different choices for α. The rightmost bin consists solely of SNPs perfectly correlated with the focal SNP. (b) The mean number of family positives in each *r* bin in families with a true positive. (c) The percentage of SNPs in each *r* bin. Probabilities and ratios plotted in log10 scale.

The above approaches for maximizing the true positive rate are based solely on significance testing. As an alternative, we explored using the rank of P-values within a SNP LD family to maximize true positives. For full-rank analyses, the focal (i.e., effect) SNP had the lowest P-value in 77% of the families; in 86% the focal SNP had one of the two lowest P-values. Figure 7a shows the proportion of all families for which the focal SNP in fact has the best P-value among all members of its family as a function of *k*. The probability that this fortunate situation occurs increases markedly with *k* to nearly 80% at the full dimensionality simulated. The situation is even more favorable if just the families with significant SNPs are included in the analysis.

**Figure 7.**
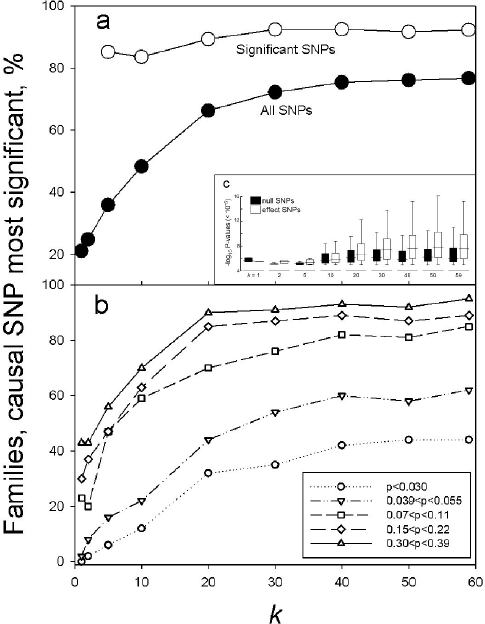
Percentage of families where the focal SNP has the best P-value, as a function of tested dimension. (a) Results for all SNPs (filled circles) and just those families where at least one SNP is significant at P<10^−5^ (open circles). (b) Results for 5 representative deciles of focal allele frequencies (p). The above results clearly show that relying only on statistical testing in the presence of LD will produce high family-discovery rates. (c) Distribution of logs of inverse significant P-values (α=10^−5^) for effect and null SNPs as a function of the number of measured traits, pooled over families. Boxes: IQR and median; whiskers: non-outlier range.

Analysis of the DGRP data shows that a much larger number of SNPs are highly correlated with rare minor alleles compared with common minor alleles. We therefore expected that the sites with low allele frequencies would be much less likely to have the focal effect SNP with the lowest P-value. Figure 7b shows that this is indeed the case. For these parameter values, causal SNPs are very likely to be correctly identified with high-dimensional analyses in families with allele frequencies over 0.07.

More generally, the results in Fig. 7 are driven by greater dissimilarity between the distributions of P-values for correlated effect and null SNPs at increasing *k* (Fig. 7c). Both distributions become increasingly biased toward lower P-values as *k* approaches *d*, but P-values of effect SNPs are improved more than those of null SNPs. This discrepancy is likely a general outcome of increasing statistical power.

These results indicate that when power, effect size and minor allele frequency are large enough, it can be effective to test all SNPs for statistical significance, and then compare those results within LD families. For cases where there is likely to be lower power than we have simulated, the probability that the most extreme P-value correspond to effect SNPs will certainly be lower than in our simulations. However, retention of a sample of the SNPs with the best P-values within their families would remain an effective strategy for winnowing sites to validate.

## Discussion

Our simulation results have shown several advantages of multivariate analyses for association studies that are likely to be general. Power generally rises and false discovery rate falls with the number of phenotypes that are measured. In addition, the ability to discriminate between true positives and positives caused by linkage disequilibrium with a causal SNP also increases with the number of traits measured. We also show that association analyses that explicitly take into account the SNPs that are highly correlated with significant SNPs will be superior to those that do not.

Despite the fact that we based our simulations on the allele frequency and linkage disequilibrium (LD) distributions in a specific data set, the Drosophila Genome Reference Panel (Mackay et al. 2012), our results address general features of multivariate analyses. The distribution of allele frequencies in the DGRP is a typical U-shaped distribution. Mackay et al. (2012) showed that the influence of LD in the DGRP was less than in other metazoan populations, but substantial LD is found even in this population, as our unpublished analyses show. By focusing our LD simulations on SNPs that are in LD with other SNPs, we capture the effects of LD with respect to a focal SNP, which will be similar in any population.

Our first major result is that power in GWAS increases dramatically with the number of measured traits (*k*), up until the number of traits measured captures the true dimensionality of the phenotype, *k*=*d*. Similarly, false discovery rate falls under the same assumptions. These qualitative results hold over a wide range of power, and choice of significance thresholds, as shown in Figs. 1–3. The gains can be very dramatic: for example, in the case of maximum true dimensionality in Fig. 3, power increases by more than 10-fold when the number of traits increases from one to the maximum number of traits, while the false discovery rate falls by 3-fold. The likely cause of this is that the directions of simulated effects are most distinct in the space that they actually reside in. Effects that are in very different directions in a high-dimensional space will often have very similar effects in a low-dimensional subspace. On the other hand, increasing the number of measured traits *k* above true dimensionality *d*, simply adds error due to non-genetic effects without adding new information. Consequently, when the number of measured traits exceeds true dimensionality, power falls and the false discovery rate increases. However, the rate of gain for measuring an additional trait is greater below the correct dimensionality than the rate of loss above the correct dimensionality. For example, for the parameters in Fig. 3, if the true dimensionality is 20, increasing the measured traits from 1 to 20 increases power by a factor of 8, while measuring an extra 39 traits, to *k*=59, decreases power by only one-fourth.

Another aspect of our results that requires some comment is the correlation between power and true dimensionality (*d*) when the number of traits analyzed (*k*) is also allowed to increase. This is equivalent to moving up the diagonal of Figures 1 and 2. Our simulation algorithm holds the total amount of genetic variation across all *d* dimensions constant, so that in the *d*=1 case all genetic variation is concentrated along just one vector. The gain in power with *d* is best explained by the spreading of effect vectors into higher dimensional spaces, so that their directions become increasingly distinct. At *d*=1, all the genetic variation that is not explained by the focal SNP is part of the error variation against which that effect must be detected. In higher dimensions, only SNPs whose effects are in similar directions to that of the focal SNP contribute to the background unexplained variation. For a particular experimental population, *d* is not under the control of the experimenter, who can only choose *k*.

Although our results may be new to some within the community doing GWAS, they are largely expected based on straightforward geometric and statistical arguments. Multivariate methods take advantage of the entire space of possible effects for testing. If we measure *k* traits, we could do *k* univariate tests, which will be successful at detecting effects that are large in the direction of each measured trait. However, if a genetic effect of the same relative magnitude is spread over a large proportion of the measured traits, it may be undetectable to univariate analyses, but a multivariate analysis will be just as likely to detect it as if the entire effect were on a single trait. When we do not measure all the trait effects that a SNP produces we are measuring *projections* of true genetic effects, and these can be very misleading.

It is important to realize that we have examined the average power over all the SNPs that explain our simulated genetic effects. O’Reilly et al. (2012) performed a complementary set of simulations to ours, in which they considered the power to detect individual SNP effects against varying backgrounds of more or less correlated residual variation. As in our simulations, they found that the power of a multivariate approach was generally much greater than univariate analyses, but that there were particular orientations of SNP effects relative to the residual variation where univariate analyses performed slightly better than a multivariate one. Such cases undoubtedly occur in our simulations for individual SNPs whose orientations are, for example, exactly in the same direction as a measured trait (or an eigenvector). However, we sampled SNP effects randomly from throughout the multivariate space simulated, and our results reflect the average power to detect the entire set of SNPs, not individual cases.

We have also shown that increasing the number of dimensions analyzed can greatly aid in distinguishing true positives and SNPs that are significant because of linkage (gametic-phase) disequilibrium (LD, Fig. 7) with a causal SNP. Presumably the mechanism that produces this effect is similar to those that increase power and reduce FDR in the absence of LD– more information is added with every trait measured. SNPs with no real effects may happen to diverge from the null in any one dimension, but are increasingly unlikely to do so when averaged over all dimensions.

The asymmetry in the disadvantages of measuring too many vs. too few traits depends on some important but biologically and operationally justifiable assumptions about the nature of the traits measured. We have defined traits based on a principal components rotation of the genetic variance-covariance matrix. This defines our trait 1 as the combination of the measured phenotypes with the most genetic variation, trait 2 as the combination with the second most variation, etc. This mimics experiments in which the full set of traits is measured before analysis, so that intelligent decisions can be made about which combinations of traits are most informative. This is a common approach to the analysis of multivariate data sets [e.g., 3], that ensures both that no two variables explain the same variance (i.e., variables are conditionally independent), and that variables, when considered jointly, account for the entire variance in the data. The use of a principal components definition of traits explains the asymmetry noted above. In making the transition from measuring *k* traits vs. *k*+1 informative traits, we have always chosen to add the most informative trait. Similarly, when going beyond the true dimensionality, the *d*+1*th* contributes less variation than the *dth* trait, so less misleading error variance is contributed with each extra trait.

This situation may be rather different when considering whether to measure and analyze an additional trait whose variational properties are not known. Here, one can imagine different outcomes. For morphological traits, the majority of variation is frequently in overall size, so measuring any one part of the body is likely to capture a great deal of size variation. In this case the first trait measured is likely to capture more variation than the second, just as with the principal components approach. On the other hand, if mixtures of different types of traits are studied, for example life history, physiological and behavioral traits, it is very possible that the first trait measured will be less informative than a subsequent trait. Given these considerations, we recommend that wherever possible a preliminary assessment of the true genetic dimensionality of the suite of traits under study be undertaken prior to an association study. Several approaches have been utilized to estimate trait dimensionality [26–30], but their implementation in genomic analysis must become widespread if we are to assess the true statistical power of GWAS. Ultimately, the decision whether to include an additional trait in an analysis depends on the amount of new genetic variation brought by the trait, which can be estimated as its heritability after partialling out the effect of all other measured traits. PCA and other eigendecomposition methods are computationally inexpensive procedures to generate sets of conditionally independent traits. Multiple regression of new on previously measured traits can help determine whether additional phenotypes significantly expand the dimensionality of a data set.

A major challenge to all real genome-wide association studies is the presence of linkage disequilibrium, LD. When the genotypes at different SNPs are correlated, a significant association at one will predict that a significant association will be found for the other. When we simulated such cases, we found that, as expected, SNPs with real effects tended to have lower P-values than those without effects. The key result however, is that the tendency of P-values to distinguish true from LD-correlated effects was greatly enhanced by increasing the dimensionality of the analysis. This may be a general side-effect of the increasing statistical power [31,32], which suggests an avenue to seek high-power conditions that ensure robustness of GWAS to the violations of the assumption of independence among tested SNPs.

We caution that the actual power and false discovery rates we find in our simulations are dependent on a large number of assumptions built into our simulations. Heritability of the traits considered here (Fig. S4), which determines the upper boundaries of wing shape dimensionality, determines the precise extent whereby matching sampling (*k*) and true (*d*) dimension translates into a power increase. Highly integrated phenotypes where a few dimensions explain most of the genetic variance will see a diminished advantage in adding new traits relative to our study, whereas complex phenotypes comprising traits from multiple systems could see even stronger gains in power at higher dimensions. No general conclusions can be drawn from the numerical values in our results. What we believe can be generalized from our results are these overall trends: given the high dimensionality of most, if not all, phenotypes, measuring and analyzing more traits is better than measuring fewer.

## Conclusions

Despite the statistical advantages to multivariate analyses that we have demonstrated, the decision to embark on such a study depends on the nature of the question under study, and the costs of doing so. In many cases, the principal drawback of doing a multivariate association study is the cost involved in measuring more traits, which is compounded by the costs of dealing with additional factors such as population structure [e.g., 33] and cross-tissue variance [34,35] in a multivariate context. Multivariate studies will be greatly facilitated by the development of high-throughput phenomic techniques [16,36].

Traditional power analysis focuses on magnitudes of effects and sample sizes, but our results strongly suggest that the number of traits analyzed in relation to the true dimensionality also greatly affects the expected power of a GWAS. We recommend that explicit analyses of phenotypic dimensionality be undertaken as part of the designing a GWAS, so that an informed balance between the number of traits and the number of genotypes to assess can be achieved. This is likely to be especially critical in genomic studies like eQTL analyses of transcript abundances [37,38], where the cost of generating and sequencing additional genotypes far exceeds those of acquiring nearly complete phenotypic information.

It is clear that the true dimensionality of complex phenotypes is generally larger than one [16]. Consequently, we recommend that multivariate approaches be adopted much more widely. This will result in gains in power and decreases in false discovery rates, particularly in the presence of linkage disequilibrium. We can also anticipate that when the directions of effects are included in the interpretation of SNP effects, multivariate approaches will increase the interpretability of the mechanisms of genetic effects, and the complexity of the pathways by which they do so. All these factors suggest that identification of causal variation will become easier as truly phenomic approaches to GWAS become more widespread.

## Acknowledgements

We thank the FSU Research Computing Center for use of their computational resources. We thank Ian Dworkin and two anonymous reviewers for comments. Phenotypic data used as the basis for our simulation parameters was collected by Jessica Nye.

